# EthSEQ: ethnicity annotation from whole exome sequencing data

**DOI:** 10.1101/085837

**Authors:** Alessandro Romanel, Tuo Zhang, Olivier Elemento, Francesca Demichelis

## Abstract

Whole exome sequencing (WES) is widely utilized both in translational cancer genomics studies and in the setting of precision medicine. Stratification of individual's ethnicity is fundamental for the correct interpretation of personal genomic variation impact. We implemented EthSEQ to provide reliable and rapid ethnicity annotation from whole exome sequencing individual's data and validated it on 1,000 Genome Project and TCGA data demonstrating high precision (>99%). EthSEQ can be integrated into any WES based processing pipeline and exploits multi-core capabilities. Source code, manual and other data is available at http://demichelislab.unitn.it/EthSEQ.

## Introduction

Interrogation of the entire coding genome for germline and somatic variations through Whole Exome Sequencing (WES) is rapidly becoming a preferred approach for the exploration of large cohorts (such as The Cancer Genome Atlas initiative) especially in the context of precision medicine programs (Beltran *et al.*, 2015). In this setting, the estimation of individual's ethnical background is fundamental for the correct interpretation of variant association studies and of personal genomic variations importance (Zhang *et al.*, 2016; Petrovski and Goldstein, 2016; Spratt *et al.*, 2016; Price *et al.*, 2006). To enable effective annotation of individual's ethnicity and improve downstream analysis and interpretation of germline and somatic variations, we developed EthSEQ, a tool that implements a rapid and reliable pipeline for ethnicity annotation from WES data. The tool can be used to annotate ethnicity of individuals with germline WES data available and can be integrated in any WES-based processing pipeline. EthSEQ also exploits multi-core technologies when available.

## Implementation

EthSEQ provides an automated pipeline, implemented in R language, to annotate the ethnicity of individuals from WES data inspecting differential SNPs genotype profiles while exploiting variants covered by the specific assay. The tool requires as input genotype data at SNPs positions for a set of individuals with known ethnicity (the *reference model*) and a list of BAM files representing a multi-individual WES cohort. EthSEQ then infers and annotates the ethnicity of each individual using an automated procedure (Figure 1a). EthSEQ returns a table with detailed information about individual's inferred ethnicity, including aggregated and sample-based visual reports.

**Figure 1:**
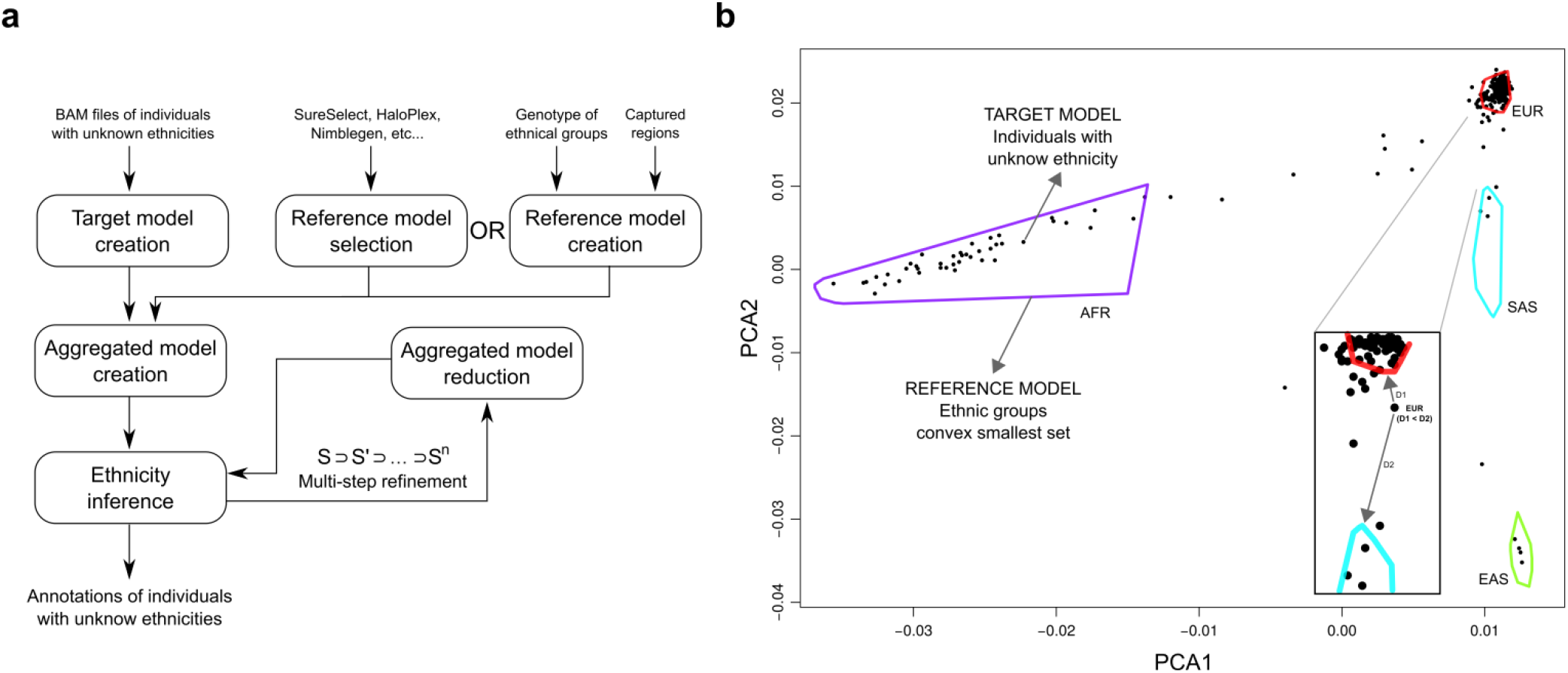
EthSEQ analysis pipeline: a) Illustration of the pipeline implemented by EthSEQ to infer ethnicity of individuals with germline WES data available. Both standard and multi-step refinement procedures are shown. b) Example of EthSEQ visual report representing the 2-dimensional space built with the first two PCA components. Polygons represent reference model ethnical groups, while black points represent individuals with unknown ethnicity. Inset shows how distance from reference ethnical groups is used to infer ethnicity when an individual lies outside polygons.

The *reference model* builds on genotype data of individuals with known ethnicity, such as the 1,000 Genome Project individuals, which is used here to construct platform-specific reference models relying on the most conserved ethnic groups EUR (Caucasian), AFR (African), EAS (East Asian) and SAS (South Asian) for multiple WES designs: Agilent HaloPlex, Agilent SureSelect and Roche Nimblegen (models are available online, **Supplementary Methods**). More generally, given a description of WES captured regions (in BED format) and genotype data files (PED or VCF format) of a set of individuals annotated for ethnicity, a procedure to automatically generate a reference model is also provided by EthSEQ.

The *target model* is then created from the input list of individual's germline BAM files that are genotyped at all reference model's positions using the genotyping module offered by ASEQ (Romanel *et al.*, 2015), hence exploiting multi-core capabilities. Depth of coverage >=10X and read/base mapping qualities >=20 are required by default to guarantee confident genotype calls (Romanel *et al.*, 2015) (parameters can be tuned by the user). Principal component analysis (PCA) is then performed by means of *smartpca* module (Price *et al.*, 2006) on aggregated target and reference models genotype data considering only SNPs with appropriate overall call rate (default 100%, tunable by the user). The 2-dimentional space defined by the first two PCA components is then automatically inspected to, first, generate the smallest convex sets identifying the ethnic groups described in the reference model and then to annotate the ethnicity of the individuals of interest (Figure 1b). Individuals positioned inside an ethnic group set (or intersecting group sets) are annotated with the corresponding ethnicity and with the INSIDE label; individuals lying outside ethnic group sets are annotated with the nearest (Euclidean distance) ethnic group and CLOSEST label (Figure 1b, **inset**).

To better discern ethnicity annotations across ancestrally close groups within a study cohort (for example Ashkenazi and Caucasians), a multi-step inference procedure is provided. Given sets of ethnic groups 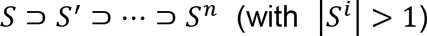 ethnicity of each individual is first inferred using all ethnic groups in S. Both reference and target models are then reduced to S' individuals only and the analysis is reiterated. The refinement procedure is repeatedly applied till converging to ethnic groups in S^n^. Finally, individuals ethnic annotations is compiled by recursively combining all refinement steps results.

## Performances and Results

To test the performances of EthSEQ ethnicity inference method we first considered 1,000 Genomes Project individuals genotype data (N=2504, release 20130502). We randomly divided those individuals into reference and target model groups while preserving the ethnic groups proportions, and ran EthSEQ relying only on SNPs in WES platform-specific captured regions (**Supplementary Methods**). Individuals’ ethnicities were all correctly classified (100% precision). More than 97% of the individuals were annotated with INSIDE label, overall demonstrating strong confidence of the calls (**Table S1**). Further, germline WES data (Agilent SureSelect v2 assay) from TCGA PRAD individuals was considered (The Cancer Genome Atlas Research Network, 2015). Of the 333 individuals in the dataset 191 had reported interview-based ethnicity classification categorized into *White* (N=163), *Black or African American* (N=22) and *Asian* (N=6) populations. 160 *White* individuals were annotated by EthSEQ as EUR, 21 *Black or African American* individuals as AFR and 4 Asian individuals as EAS or SAS. Of the six discordant annotations (**Table S2**) three demonstrate genetic profiles that are compatible with admixed ethnical background; the remaining three suggest potential reporting mistakes in interview-based annotations. Of the 142 individuals with missing original ethnicity information 122 had high confident EthSEQ annotations (INSIDE label), overall providing a complete and robust classification for TCGA PRAD dataset (Figure 1b, **Figure S1, Table S3**). Notably, starting from the list of 333 germline BAM sample's files and using our pre-built reference model for Agilent Sure Select v2, the overall pipeline computation required less than 12 hours to complete (~2 minutes per sample), exploiting 30 cores of the in-house machine (4 Intel® Xeon CPUs E7540 at 2GHz with 12 cores in hyper-threading mode) for the target model creation.

The effectiveness of the multi-step refinement analysis was recently proven in a precision medicine setting study (Zhang *et al.*, 2016) were ethnicity based stratification was key to interpret the relevance of germline cancer-associated variants. Specifically, our analysis ruled out the possibility that the high fraction of germline cancer-associated variants observed in the clinical cohort of 155 patients with metastatic tumors (Beltran *et al.*, 2015) was due to the presence of Ashkenazi inheritance (Carmi *et al.*, 2014) shown to carry high percentage of cancer-associated variants. Provided an Agilent HaloPlex reference model including Ashkenazi genome data (Carmi *et al.*, 2014) the identification of Ashkenazi individuals required the multi-step analysis to precisely discern them from the ancestrally close European individuals; 34.8% of Ashkenazi individuals were identified confirming the anticipated fraction of about 30% based on an internal cancer registry.

To measure the impact of SNPs availability on precision, we first extended the performance analysis we performed on the 1,000 Genome Project data by randomly down-sampling the number of SNPs used by EthSEQ (**Supplementary Methods**). **Figure S2a** shows that by reducing the number of variants of 2 orders of magnitudes, precision is greater than 99%. Similarly, 1,000 SNPs on a TCGA dataset (**Figure S2b**, **Supplementary Methods**) result in precision >97%. Of note, when reference models including ancestrally close ethnic groups are used (e.g. Ashkenazi and Europeans individuals), the multi-step inference procedure guarantees high sensitive calls also in the presence of low SNPs availability (**Figure S3**). Overall, this data indicate that EthSEQ is also amenable to targeted sequencing NGS data.

## References

Beltran, H. et al. (2015) Whole-Exome Sequencing of Metastatic Cancer and Biomarkers of Treatment Response. JAMA Oncol., 1, 466–474.

Carmi, S. et al. (2014) Sequencing an Ashkenazi reference panel supports population-targeted personal genomics and illuminates Jewish and European origins. Nat. Commun., 5, 4835.

Petrovski, S. and Goldstein, D.B. (2016) Unequal representation of genetic variation across ancestry groups creates healthcare inequality in the application of precision medicine. Genome Biol., 17.

Price, A.L. et al. (2006) Principal components analysis corrects for stratification in genome-wide association studies. Nat. Genet., 38, 904–909.

Romanel, A. et al. (2015) ASEQ: fast allele-specific studies from next-generation sequencing data. BMC Med. Genomics, 8, 9.

Spratt, D.E. et al. (2016) Racial/Ethnic Disparities in Genomic Sequencing. JAMA Oncol., 2, 1070.

The Cancer Genome Atlas Research Network (2015) The Molecular Taxonomy of Primary Prostate Cancer. Cell, 163, 1011–1025.

Zhang, T. et al. (2016) Germline Variants and Secondary Findings in a Cancer Precision Medicine Cohort. In, LABORATORY INVESTIGATION, p. 461A–461A.

